# Reduced nutritional quality of plants due to climate change

**DOI:** 10.64898/2026.04.08.717215

**Authors:** Fabio Berzaghi, David Makowski

**Affiliations:** Laboratory of Climate and Environmental Sciences (LSCE) - UMR CEA/CNRS/UVSQ; Gif-sur-Yvette, France; Sasakawa Global Ocean Institute, World Maritime University, Fiskehamnsgatan 1, 211 18 Malmö, Sweden; INRAE AgroParisTech Université Paris-Saclay, Unit Applied Mathematics and Computer Sciences, 91120 Palaiseau, France

## Abstract

The effect of climate change on the nutritional quality of plants is poorly understood even though it may have major implications for the food chain and ecosystems. Here we use a dataset covering > 1450 plant species to identify the factors driving plants nutritional properties and develop global projections. We reveal that plant type, CO_2_, and solar radiation are major drivers controlling nutritional properties. Projections for 2050 show a decline in nutritional quality (-8%, on average at the global scale), measured as the protein to fiber ratio, a strong decrease in minerals (-18%), and a small decrease in digestibility (-3%). Plants in arid and tropical areas will experience the largest decline in quality, which will decline minimally in temperate areas and improve in cold, and polar regions. Quality trends will be opposite in grasses. These results have important implications for livestock management and wildlife conservation.

## Main Text

Forage plants are an important source of nutrients for livestock, wild animals, and humans. Plants contain different amounts of protein, fibers, and minerals, which vary across plant types and are influenced by environmental conditions^4–6^. Within plant types, there are differences in protein, fiber, and minerals content^4^. The ratio between proteins and fibers decreases during growth but is also affected by temperature and moisture^6^. This ratio determines feed/food quality, in addition to its minerals content. Plants high in protein and low in fiber are considered of high nutritional quality also because they are more digestible (proportion of digestible biomass) than plants with higher fiber content^6,7^. Feed quality is positively linked to reproductive success, weight gain, milk and meat production from livestock and can affect animal behavior and methane emissions^4,5,7,8^. Nutritional values are also critical for wild animals who cannot easily switch diet during periods of low-quality feed or rely on supplements, as compared to domestic animals^9^. Lastly, human nutrition is directly and indirectly dependent on the quality of plants (especially, on protein content) used as food or as feed for animals^1^.

The ongoing changes in weather and climate are affecting the nutritional quality of both cultivated and wild plants^1,2,10,11^. However, most of the research effort has focused on the effects of climate on crop yield or on habitat suitability^3,12–14^. In previous experimental studies, effects on nutritive properties have been evaluated mostly at local scales, often using one or two treatments (elevated CO_2_, temperature, drought, etc.)^1,15^. These experiments cannot fully reproduce the complexity of climate in which factors interplay. For example, the projected increase in global temperature is also associated with increased precipitation and modified seasonality^14^ with uncertain eco-physiological responses of plants. Recent global modeling efforts showed a global reduction in protein and minerals content of crops by 2050 although with considerable regional variability^16,17^.

The majority of these previous studies were conducted on model plants or cereals, which represent only one plant type among many others. Conversely, research on the effects of climate change on the nutritional quality of other plant types and wild plants is very limited^2,10^. Wild plants are in some ways more vulnerable to global changes than crops because they cannot benefit from adaptive strategies implemented by farmers, such as fertilization, irrigation or improved cultivars. As herbivores are extremely dependent on the nutrient content of forage plants, future environmental changes could disrupt plant-animal interactions^18^ and make some natural regions less suitable for herbivores^5^ with unknown consequences throughout the food web. A more comprehensive assessment of changes in nutritional quality should also include fiber content and digestibility, which have so far been neglected. Because of the multiple implications of plant nutritive properties, a reliable prediction of these properties to climate change would provide valuable information for food security risk assessment and biodiversity conservation.

Here, we employ a global dataset to analyze the variation of nutritive properties – protein to fiber ratio, minerals, and digestibility measurements spanning several decades and countries – across a diversity of plant types. Based on machine learning, we identify the main factors influencing these plant nutritional properties and assess the consequences of climate change on plant quality by factoring for plant types, climate zones, and land cover. Finally, we generate global surface maps of nutritional qualities under both current and future climate conditions. Our assessment will contribute to inform land management, conservation actions, and policy making.

## Results

The data included a total > 9000 records of nutritional properties over 238 unique locations which covered a broad range of climates and ecosystems (Extended Data Fig. S1). The data included 1453 species divided in five plant types: 772 trees, 173 legumes, 221 shrubs, 167 herbs, and 120 grasses.

### Relationship between nutritive properties

In our data, protein and fiber were negatively correlated and more than 10% of the variation in protein was explained by fibers (Fig. 1a, R^2^ = 0.11, p-value < 0.0001). Protein was negatively associated with fiber also within each plant type (Fig. S2, p-value < 0.0001). Digestibility was positively correlated with (r = 0.20, p-value < 0.0001) and affected by protein (Fig. 1b, R^2^ = 0.08, p-value< 0.0001). Digestibility was negatively correlated with fiber (r = -0.52, p-value< 0.0001) and decreased linearly as a function of fiber (Fig. 1c, R^2^ = 0.18, p-value< 0.0001). These two relationships were significant within all plant types, particularly as fiber explained an important portion of change in digestibility (R^2^ = 0.30-0.43, Fig. S2). The Pearson correlation between digestibility and minerals was positive (r = 0.32, p-value < 0.0001), but the regression was significant only in trees, shrubs, and herbs, but not in grasses and legumes (p-value < 0.05, Fig. S2).

**Fig. 1.**
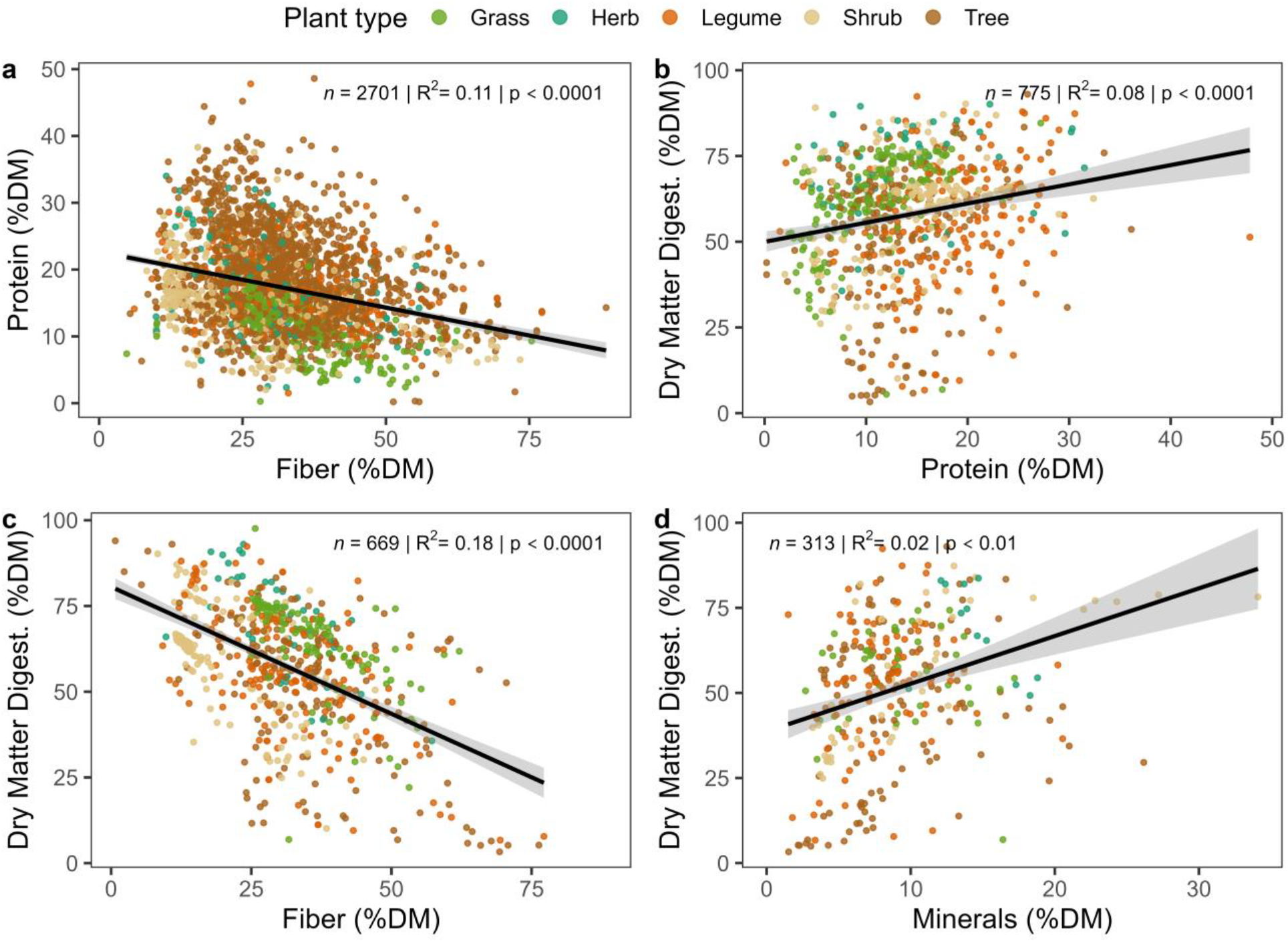
Correlations between plant nutritional properties based on measurements and linear model fit. (a) Protein as a function of fiber; dry matter digestibility as a function of (b) protein, (c) fiber, and (d) minerals. Regression lines for each plant type are shown in Extended Data Fig. S1 and complete test results are in Extended Data S1.

Analysis of differences among plant types showed that (from lower to higher average value): grasses, shrubs, herbs, and legume differed in average protein content (Sidak test, p-value < 0.0001), whereas trees had intermediate protein content (Extended Data Fig. S3a). Fibers were statistically different in shrubs, legumes/trees, and grasses (Extended Data Fig. S3b, p-value < 0.0001). Legumes, grasses, and herbs had statistically different minerals content (Extended Data Fig. S3c, p-value < 0.001); digestibility was only statistically different in herbs compared to the other groups (Extended Data Fig. S3d, p-value < 0.0001). Average plant properties, complete statistical pairwise tests, and statistical grouping are shown in Extended Data S1.

### Global drivers of plant nutritive values

We used environmental variables (mean annual temperature and precipitation, precipitation seasonality, soil type, solar radiation, CO_2_, and actual evapotranspiration) and plant type to predict nutritional properties with a Random Forest (RF) model. Based on RF, we ranked predictor variables according to their importance. The importance of each predictor is measured as the increase in the mean square error of the model resulting from a random permutation of the values of the corresponding input variable^19^. Plant type was the most important variables when predicting three out of four nutritional properties, particularly for protein and fiber (Fig. 2). Random permutation of the values of the plant type input increased the model MSE between 34% and 55% depending on which nutritional property was predicted. Temperature and CO_2_, which are the factors most studied experimentally, and solar radiation had an important influence on most properties. (Fig. 2). Soil type and precipitation were important for predicting, respectively, minerals and fibers, but less for the other properties.

**Fig. 2.**
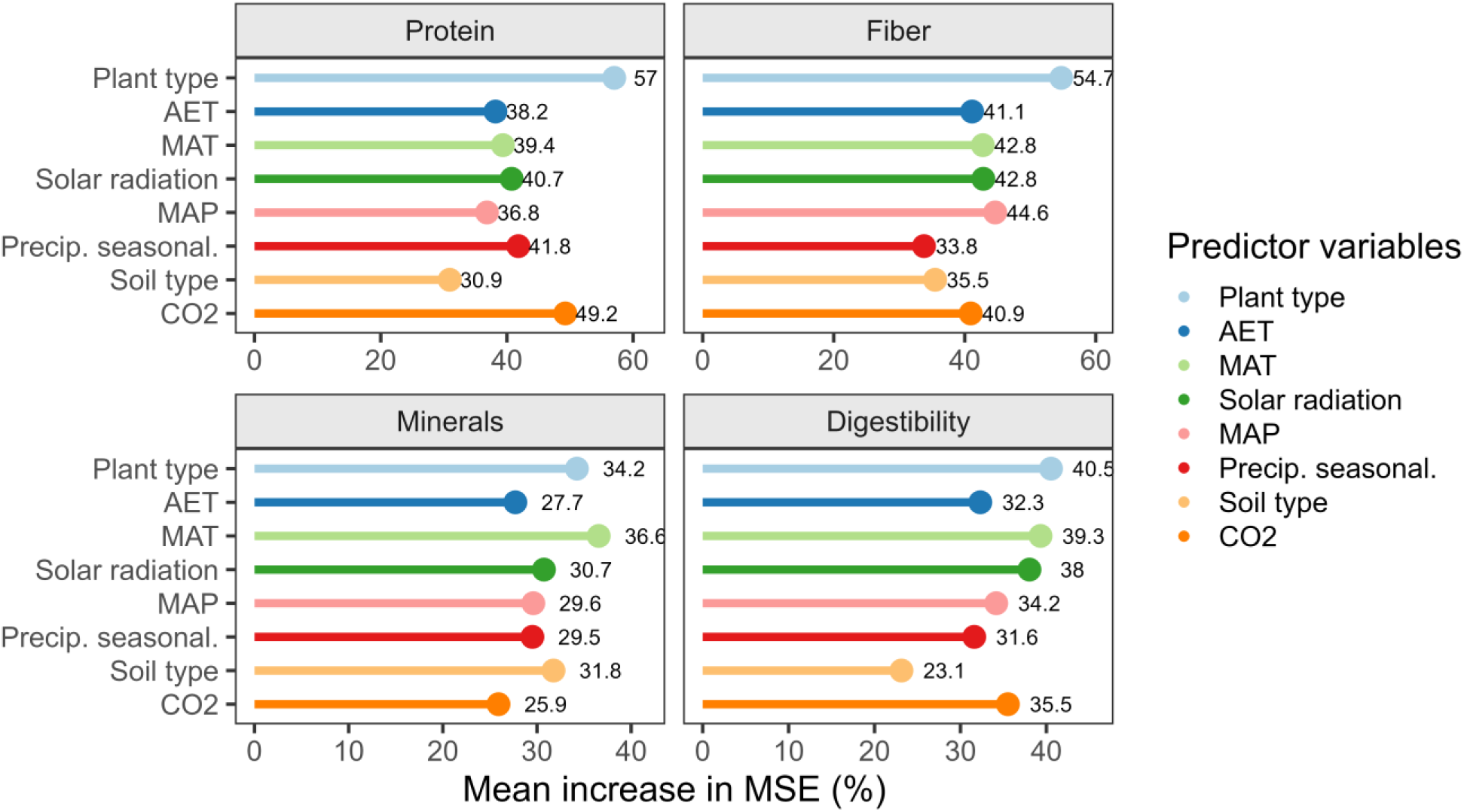
Contributions of different predictors to plant nutritive properties computed with a Random Forest model. The contribution of each predictor is measured as the increase in the mean square error of the model resulting from a random permutation of the values of the corresponding input variable.

### Global maps of nutrient properties

The Random Forest model was used to predict nutritional properties (protein, fiber, minerals, and digestibility) in 2050 according to projected changes in climate-related predictors in +2° (SSP2: Middle of the Road) and +4° (SSP4: Inequality, A Road divided) climate trajectories ^20^. Here we present the global projected map of the protein to fiber ratio (PFR), due to its importance as a proxy for nutritional quality (methods), and differences between present and future climate in PFR, minerals, and digestibility. Differences in nutritional properties across the two climate change scenarios were small (2-5%), mostly because of the relatively short time horizon considered (2050) (methods). Consequently, our results represent the average of the two scenarios.

The spatial distribution of PFR revealed that quality was lower at mid-latitudes and in subtropical areas, intermediate in tropical areas, and higher at high latitudes and some arid hot areas (Fig. 3a). These patterns result from the partially specular distribution between protein and fiber (Extended Data Fig. s4 and s5). The range of empirical data across climates was larger than the range predicted by the RF model (Fig. 3b). However, average predictions from the model were close to empirical averages (Fig. 3b and Extended Data Table s2) with the exception of polar climate where only few observations were available. Our projections suggest that by 2050 PFR will decrease globally by 8% but with high spatial variability (s.d. 12%, Fig. 3c). The largest losses of nutritional quality are projected to occur in arid climates (-13%) and tropical regions (-9%). Losses are smaller in polar (-7%), temperate (-7%), and cold areas (-3%) (Fig. 3c and 3d). Quality will increase slightly in some mid-and high latitude zones, and a few sub-tropical areas (Fig. 3b and 3c).

**Fig. 3.**
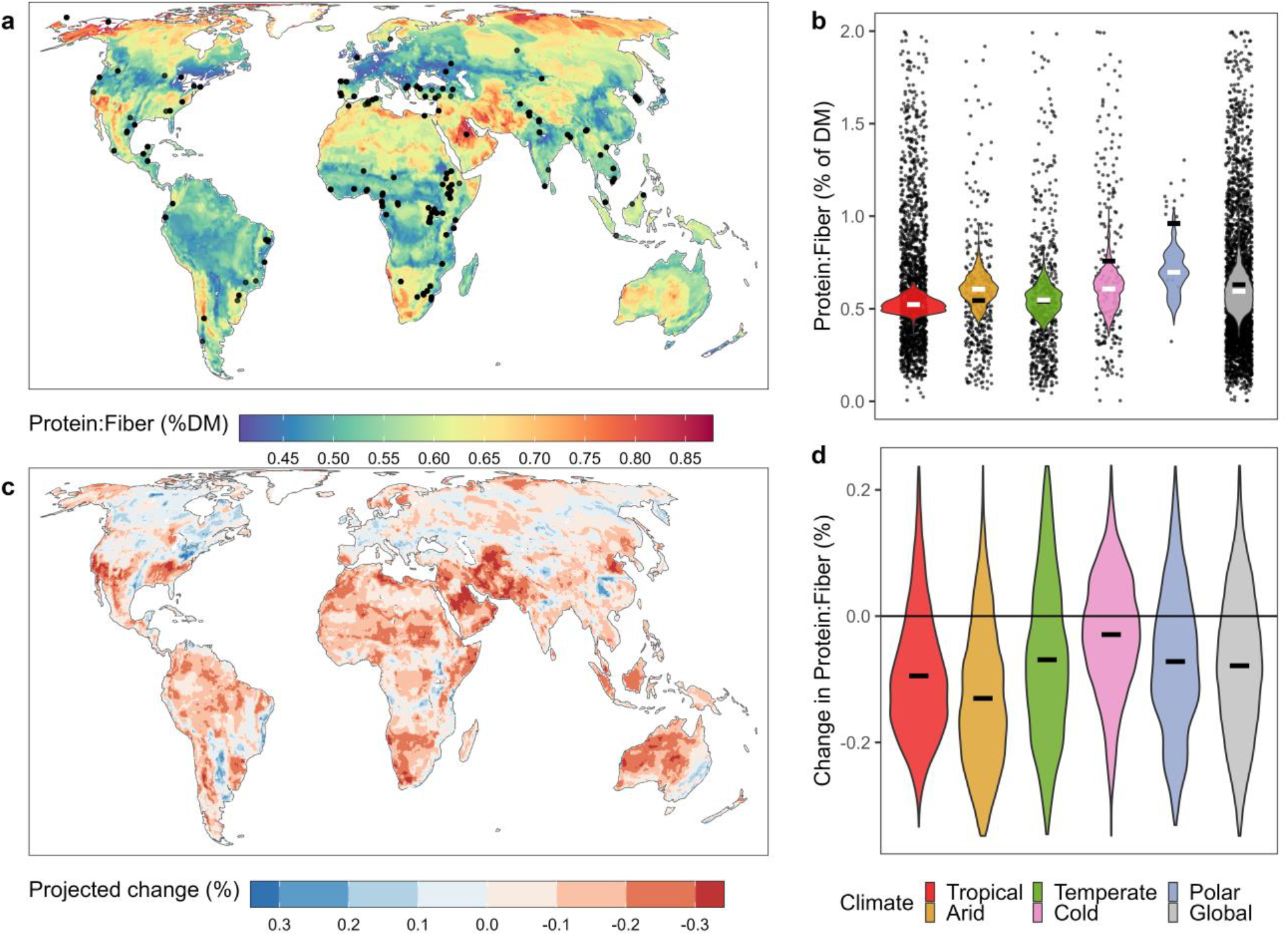
Global patterns of observed and predicted protein to fiber ratio. Predicted global map of protein to fiber ratio in 1961-2018 (a) and predicted (violin charts) vs. observed values (points) for the same period (b), global map of projected changes of protein to fiber ratio by 2050 compared to 1961-2018 (c), and ranges of predicted values (violin charts) across climate zones. Distribution of climate zones is shown in Extended Data Fig. S9. Points in (a) indicate locations where one or more plant samples were collected in their natural growing environment. The horizontal bars in (b) and (d) represent the average of predicted (black) and observed values (white).

Changes in plant PFR across continents showed clear patterns that reflect climate zones within each country and continent, but the spatial variability of change in PFR was high within countries (Fig. 4). Africa and Asia were the continents with the biggest projected decrease in plant PFR (-3-20%) because most of their countries are in arid and tropical climates. Countries in the Middle East and Southeast Asia had high projected reduction in PFR (-15-24%), including highly biodiverse and human-populated countries such as Indonesia, Pakistan, and Philippines. European countries showed both positive and negative changes in PFR; losses were observed primarily in northern countries with the exceptions of Portugal and Spain. In the Americas, some countries showed large variations, due to the wide range of within-country environmental conditions. Canada, Honduras, and Dominican Republic were the only large countries in the Americas with an overall positive change in quality (Fig. 4).

**Fig. 4.**
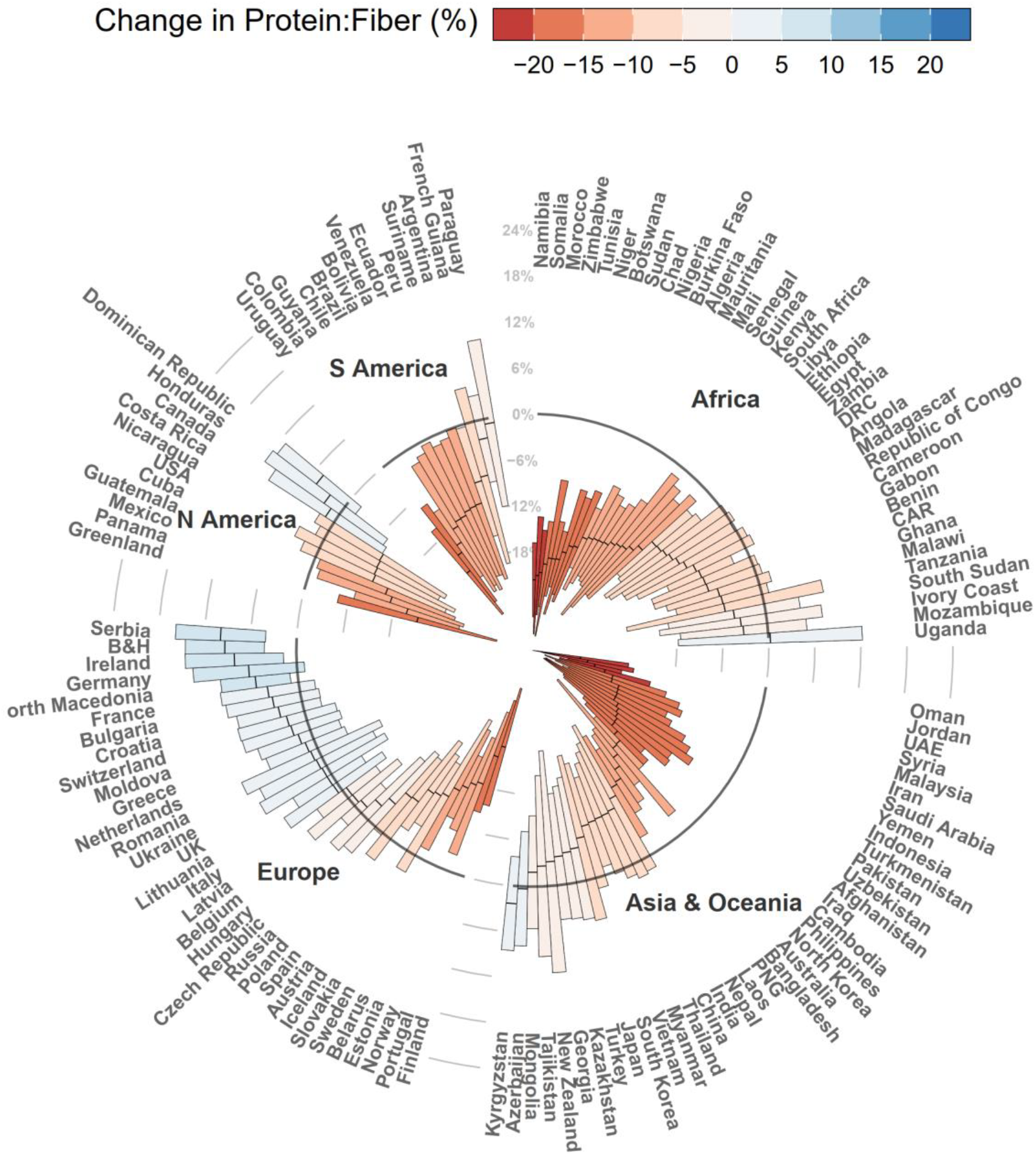
Projected changes in protein to fiber ratio quality within countries by 2050. The bold circular line represents no change (0%) and the black centerlines indicate the average value for each bar plot. The top and bottom of the bars represent 1 ± standard deviation. Countries with small areas are not shown to improve readability and can be found in the supplementary.

### Changes in nutritional properties across plant types and habitats

Plant types showed different responses to climate change in their projected PFR, although with a high degree of variability (s.d. 11-13%, Fig. 5a). Trees showed a global 1% increase in PFR, whereas PFR decreased in herbs, shrubs, legumes between 2-3%, and grasses changed was zero (Fig. 5a). These global trends masked subtle differences that appeared by analyzing future changes in natural and artificial habitats (arable land, pasture, and plantations). The global trend in artificial habitats was mostly an increase in PFR in grasses and trees and minimal changes in the other plant types (Fig. 5b). In natural habitats the trend was predominantly negative but always within a few percentage points (Fig. 5b). These results suggest that PFR could slightly improve for plants in areas currently covered by croplands. Contrarily, wild animals in natural areas might be affected by the reduction in PFR. The analysis of changes in quality in different climate zones provided additional insights (Fig. 5c). Plants in tropical and arid climates showed a consistent decrease in PFR, with the exception of grasses in arid zones. In temperate climates, which are areas heavily utilized by humans, losses were more moderate (1-3%, Fig. 5c) with the exception of grasses (+5%). In cold and polar climates, projections suggested an overall increase in PFR with the exception of grasses and herbs (only in polar). These high gains in PFR might be linked to colder climates becoming milder. These results, based on present-day land cover maps, suggest that expansion of agriculture in natural habitats might reduce nutritional quality of crops and that colder climates might become more suitable for woody plants and legumes.

**Fig. 5.**
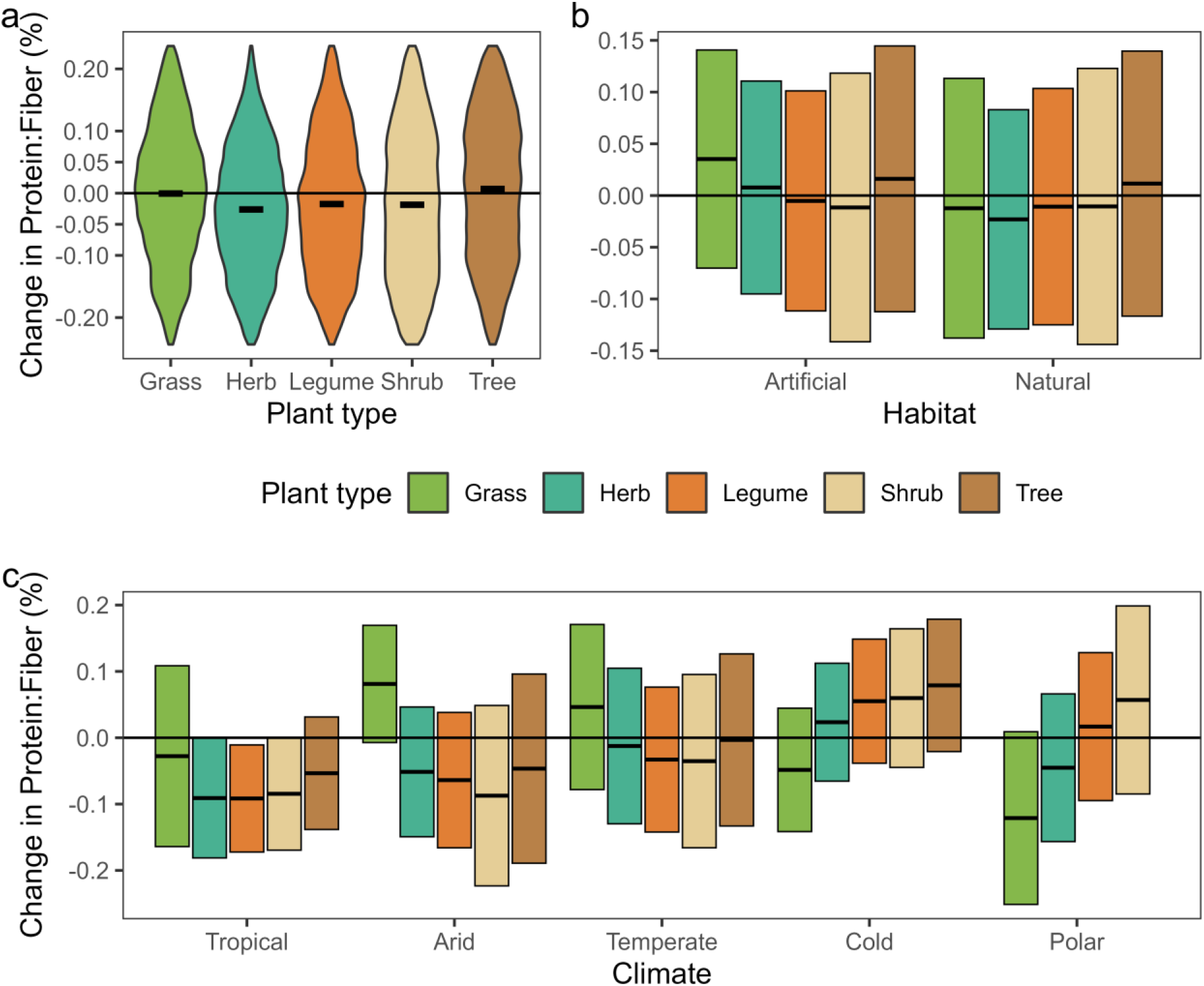
Projected changes in protein to fiber ratio in plant types in different habitats and climate regions by 2050. The black horizontal centerlines indicate the average value for each bar or violin plot. The top and bottom of the bars represent 1 ± standard deviation.

### Minerals and digestibility

Changes in PFR need to be evaluated in conjunction with changes in minerals and digestibility. Contrarily to changes in PFR which were highly variable, minerals were projected to consistently diminish between 10 to 20% and up to 30% in some areas (Extended Data Fig. S6). The trend of change in minerals across climates was opposite of what we observed in PFR. Plants in tropical areas showed the smallest decrease in minerals, followed by temperate, arid, cold, and polar climates (Extended Data Fig. S7). Differences among climates were much larger than differences among plant types, which showed very similar losses in minerals (Extended Data Fig. S7). Minerals decreased more in artificial compared to natural habitats (Extended Data Fig. S8). Losses in digestibility were limited to a few percentage points, although with large spatial variability (Extended Data Fig. S9). Arid climates showed the biggest decrease in digestibility followed by tropical and temperate areas (Fig. 6). Future changes in digestibility also seemed very similar across plant types with a general increase of a few percentage points only in some cold and polar climates (Fig. 6). Changes of digestibility in plant types within artificial and natural habitats showed similar patterns (Extended Data Fig. S8). However, digestibility in artificial habitats decreased slightly less than in natural ones.

## Discussion

Many factors influence plant nutritional properties and the relationships between them. Our analysis confirms that the negative correlation between protein and fiber, and between fiber and digestibility are consistent both globally and within plant types, although not as strong as shown previously with less species^4^. All environmental variables included in our models contributed in predicting the variability of nutritional properties. However, taxonomy, temperature, and radiation were consistently important factor in such predictions. Significant differences in measured protein and fiber between plant types and their projected changes suggest that crop species might not be representative of all plants’ responses to climate changes. The tight relationship between protein, fiber, and digestibility and the high spatial variability in PFR suggest that more accurate assessments of future changes on plant nutritional quality need to include all these properties in addition to plant type and control for solar radiation. Our results depict a nuanced picture of changes in future plant nutritional quality and, in some areas, are in contradiction with previous results that showed primarily a decrease in protein^1,15^, although others suggested that forage quality could also increase^21^. The variability in protein is primarily affected by plant type, and secondarily by CO_2_, but other factors come into play (Fig. 2). Some of these factors are rarely considered in experimental studies. Direct comparisons are difficult to make as most studies have focused on seeds of cereals and legumes. Nonetheless, there are also areas of agreement such as the global reduction in minerals^11,17^. Single-treatment experiments help assess changes in plant physiology, but these experiments are conducted in a small range of environmental conditions. We also need assessments that consider the complexity of the atmosphere and our approach captures some of these complexities by also counting for the diversity in plant types. It is thus to be expected that different types of plants respond differently in a wide range of environmental conditions that result in both positive and negative changes in nutritional quality^21^.

The ecophysiology of plants dictates that an increase in temperature and decrease in moisture triggers more investment in fibers, which decrease digestibility. As we have shown, these two properties will also change in the future with a high degree of variability and not necessarily in a negative direction. When considering daily nutritional requirements, a decrease in PFR could potentially be counterbalanced by an increase in digestibility and/or by increasing daily food intake. Increasing daily intake can be a limiting factor, particularly for animals with foregut digestion such as ruminants who need more time to process plant material compared to hindgut digesters^22^.

The potential increase in grass PFR ratio in arid areas might be relevant primarily for wild plants rather than crops because in these areas water is scarce and not all crops are adapted to these conditions. Herbivores, especially in the wild, do not consume just grass. In tropical and some arid habitats, plants are fundamental for feeding large populations of wild herbivores and some domestic ones; shrubs and trees have been studied for many years as alternative sources of fodder^23^. Strict grass consumers will be the least affected because the quality of grass could remain stable or increase, be more digestible, but will contain less mineral. Conversely, other herbivores will be negatively affected, including tropical forest-dwelling herbivores such as primates^2^, elephants^8^, and insects^24^. Future climate might also favor the growth of less palatable plants^25^, which could exacerbate the quality of feed.

We have provided the first global assessment of future changes in plant PFR and digestibility. Our results show a global decline in nutritive values but with a high degree of spatial variability and differences between plant types. Overall, plants in hot and dry climates will experience the largest decline in PFR but the smallest decline in minerals. Whereas plants in temperate and cold regions will experience small changes or potential gains in PFR coupled with unchanged or increased digestibility but the largest decrease in minerals. The net effect of these changes on wild and domestic animals’ ecophysiology and health will need to be further investigated and considered in projections and assessment risks of climate change. These changes could have repercussions for human nutrition, indirectly through changes in domestic animals’ health and through direct consumption. Even though studies have shown primarily a reduction in protein and minerals of seeds, changes in fiber and digestibility will inevitably affect the overall nutrient content of all plant organs. We have shown that including these two properties could substantially change projections of future nutritional quality. Our results have implications for global food security and nutrition, and provide guidelines for managing crops and planning conservation actions of natural ecosystems. Global-scale predictions of plant nutritional values under different climate change scenarios are critical for nature and society.

## Materials and Methods

### Leaf nutritive properties

Data of plant nutritional properties were obtained from *PNuts*, a global database of nutritional properties of plants grown in their natural environment^26^. The data contained primarily measurements of leaf nutritional properties, although some records referred to leaf and twig or stem. We selected crude protein (in the main text referred as “protein”), acid detergent fiber (“fiber”), ash (“minerals”), and dry matter digestibility (“digestibility” or DMD). These nutritional properties were the most represented in the dataset and also are the most meaningful in determining food/feed quality and palatability^7^. We complemented the PNuts data to increase taxonomic and spatial coverage through additional data found in the literature. Data included 9079 geo-located measurements spanning the period of 1961-2019. In total, the data contained 3297 records of protein, 2866 of fiber, 1954 of minerals and 962 of digestibility. Protein to fiber ratio (PFR) predictions were calculated using the random forest predictions for protein and fiber because PFR is not a directly-measured quantity.

### Climate and environmental data

We retrieved environmental and climate variables using publicly available datasets^20,27–30^. We chose environmental variables known as important drivers in plant eco-physiological processes ^1,5,7^. These variables and their sources are shown in Table S1 and were all aggregated to a 0.5° spatial resolution. Analysis in relation to habitat (i.e., land cover) were performed according to the a global map of terrestrial habitats^31^. We considered natural habitat all pixels that were covered by > 70% by one or more of the following: forest, savanna, shrubland, grassland, and wetlands. Pixels were marked as artificial habitat if they contained > 70% of artificial habitat. Desert and rocky habitat classes were not used in the land cover analysis because plant cover in these habitats might be limited.

### Nutritive metrics definitions

Protein is a crucial nutrient provided by plants that is needed for metabolic and growth activities of consumers. Protein is primarily derived from nitrogen by using a conversion factor of 6.25, although this conversion factor is probably not constant^32^. Protein content is also in influenced by physiological stage, as protein in plants declines as the plant matures at the expense of fiber which increases with growth stage^7^.

Fiber, specifically acid detergent fiber in our study, is made of cellulose, lignin, and silica^7^. As plants grow lignin and other cell constituents increase to provide structural support. Fiber is the insoluble residue after a biomass sample is treated with acid detergent to remove the majority of the other biomolecules such as protein, starch, protein, and sugars. Fiber cannot be easily digested and converted into energy; a higher fiber content reduces digestibility.

Minerals content, also referred to as “ash”, is the residue after combustion of the sample at 600° C ^7^ and includes various macro and micro nutrients (C, N, Al, Ca, Cl, Fe, K, Mg, Na, P, S, Si). Dry matter digestibility, or “digestibility” is the percentage of ingested dry plant biomass that is assimilated during digestion and used for bodily functions. Different methods are used to measure digestibility, the most common being in vitro.

All nutritional properties are expressed in percentage of dry matter (%DM).

### Statistical analysis

Observed data of nutritional properties were analyzed using linear mixed-effects models to determine the effect of plant type on plant quality, considering the study site as a random effect. We also computed global means per plant type and compared them with Tukey tests. The following formula was used for the linear mixed-effects model fitted with the lmer function of the R package “lme4”^33^: plant property∼plant type + (1|site). To study the correlation between two plant properties (X, Y) we also used a linear mixed-effects model with the following formula Y∼X+ (1|site) for the global analysis and Y∼X*plant type+ (1|site) to evaluate the effect of plant type.

### Random Forest

We trained a Random Forest algorithm to investigate the factors influencing four nutritive properties and predict their future values. The investigated properties were protein, fiber, minerals, and digestibility. We used the variables listed in Table S1 and plant type as predictors in RF to compute global maps of nutritive properties. For each property we created a global map and five separate maps for each plant type: grass, herb, legume, shrub, and tree. To create plant type-specific maps we used the global random forest regression and included in the predictors a distribution map of each plant type indicating the presence of each type. For simplicity, we assumed that each plant type could be present any location with the exception of trees that we excluded from polar climates (Fig. S6) and north of 70° north; trees can grow as far south as 55° south at Tierra del Fuego. These criteria exclude the majority of areas where trees cannot grow. Our projections assume that all plant types could be present at any location. Even though this is not universally true, aside from trees the majority of plant types are present in a large range of environments from deserts to the Arctic tundra.

Data were divided into training (75%) and test (25%) subsets. The performance of the models trained using the training dataset was evaluated by computing the root mean square error (RMSE) from the test dataset. Then, a random forest model trained with the full dataset and constructed using 600 trees, the model was evaluated by computing the percentage of variance explained from the out of bag predictions provided by the summary of a random forest model fitted to the full dataset. We reduced the effect of randomness in the predictions generated by the Random Forest model by setting the same random number generator state and we constructed 10 Random Forest models for each property. Setting the same random number generator state allows to reproduce the results. These models were then combined to generate predictions. All computations were done using the R package “randomForest” based on Breiman (2001). Overall, the predicted average was close to the empirical average (Table S2). The model did not deal well with the extreme values present in the database, which were nonetheless a small percentage of all empirical measurements. For example, only 2% of all protein measurements exceeded 25% of DM and only 2.7% were below 1% of DM.

The present-day Random Forest model for each plant property was applied to create projections on the effects of future climate scenarios based on Shared Socioeconomic Pathways scenarios SSP2: Middle of the Road and SSP4: Inequality (A Road divided). These pathways represent roughly the +2° and +4° global warming scenarios. The climate data used to create maps of future scenarios is indicated in Table S1 and the +2° and +4° were calculated from the reference preindustrial period 1850-1879^27^. In preliminary analysis we found small differences (2-5%) between nutritional properties in the +2° and +4° scenarios. We thus calculated the percentage change of nutritive properties from present-day to 2050 by averaging the results of the +2° and +4° scenarios. We preferred making projection to 2050, as opposed to 2100, because policy and management are planned on a decadal time scale at best. The +2° and +4° scenarios represent changes in global average temperature by 2100. By 2050, which was our projection target, the two scenarios will not have diverged substantially^14^, hence the small differences within our two projections. Furthermore, given the uncertainly in the actual current trajectory of global climate^14^, we estimated that presenting results as the average of two scenarios would be more appropriate.

## Acknowledgments

We would like to thank Zwanga Ratshikombo and François Bretagnolle for helping with the PNuts database, Alya Ben Abdallah for the help with setting up the initial code, and Vladimir Kazmin for providing data.

## Funding

European Union’s Horizon 2020 research and innovation program under the Marie Sklodowska-Curie grant #845265 (FB)

This work benefited from the French state aid managed by the ANR under the “Investissements d’avenir” programme with the reference ANR-16-CONV-0003 (FB)

French state aid managed by the ANR under the “Investissements d’avenir” programme with the reference ANR-11-IDEX-0004-17-EURE-0006 (FB, ABA)

## Author contributions

Conceptualization: FB

Methodology: led by FB with input from DM.

Visualization: FB

Funding acquisition: FB

Supervision: FB

Writing – original draft: FB

Writing – review & editing: all co-authors

## Data availability

All data are available from their respective sources. Unpublished data will be made available upon publication in a public repository.

## Code availability

The R code used to generate the Random Forest model and related predictions will be available upon publication at https://github.com

## Extended Data

Figs. S1 to S10

Tables S1 to S2

Data S1

## References and Notes

1. Soares, J. C., Santos, C. S., Carvalho, S. M. P., Pintado, M. M. & Vasconcelos, M. W. Preserving the nutritional quality of crop plants under a changing climate: importance and strategies. Plant Soil 443, 1–26 (2019).

2. Rothman, J. M. et al. Long-term declines in nutritional quality of tropical leaves. Ecology 96, 873–878 (2015).

3. Wang, J., Vanga, S. K., Saxena, R., Orsat, V. & Raghavan, V. Effect of Climate Change on the Yield of Cereal Crops: A Review. Climate 6, 41 (2018).

4. Lee, M. A. A global comparison of the nutritive values of forage plants grown in contrasting environments. J Plant Res 131, 641–654 (2018).

5. Lee, M. A., Davis, A. P., Chagunda, M. G. G. & Manning, P. Forage quality declines with rising temperatures, with implications for livestock production and methane emissions. Biogeosciences 14, 1403–1417 (2017).

6. Lemaire, G. & Belanger, G. Allometries in Plants as Drivers of Forage Nutritive Value: A Review. Agriculture-Basel 10, 5 (2020).

7. Burns, J. C. Advancement in Assessment and the Reassessment of the Nutritive Value of Forages. Crop Science 51, 390–402 (2011).

8. Berzaghi, F., Bretagnolle, F., Durand-Bessart, C. & Blake, S. Megaherbivores modify forest structure and increase carbon stocks through multiple pathways. Proceedings of the National Academy of Sciences 120, e2201832120 (2023).

9. Delgiudice, G. D., Mech, L. D. & Seal, U. S. Effects of Winter Undernutrition on Body Composition and Physiological Profiles of White-Tailed Deer. The Journal of Wildlife Management 54, 539–550 (1990).

10. Leffler, A. J., Becker, H. A., Kelsey, K. C., Spalinger, D. A. & Welker, J. M. Short-term effects of summer warming on caribou forage quality are mitigated by long-term warming. Ecosphere 13, e4104 (2022).

11. Myers, S. S. et al. Increasing CO2 threatens human nutrition. Nature 510, 139–142 (2014).

12. Hasegawa, T. et al. A global dataset for the projected impacts of climate change on four major crops. Sci Data 9, 58 (2022).

13. Raza, A. et al. Impact of Climate Change on Crops Adaptation and Strategies to Tackle Its Outcome: A Review. Plants 8, 34 (2019).

14. IPCC. Climate Change 2022: Impacts, Adaptation and Vulnerability. Contribution of Working Group II to the Sixth Assessment Report of the Intergovernmental Panel on Climate Change. (2022).

15. Robinson, E. A., Ryan, G. D. & Newman, J. A. A meta-analytical review of the effects of elevated CO2 on plant–arthropod interactions highlights the importance of interacting environmental and biological variables. New Phytologist 194, 321–336 (2012).

16. Asseng, S. et al. Climate change impact and adaptation for wheat protein. Global Change Biology 25, 155–173 (2019).

17. Beach, R. H. et al. Combining the effects of increased atmospheric carbon dioxide on protein, iron, and zinc availability and projected climate change on global diets: a modelling study. The Lancet Planetary Health 3, e307–e317 (2019).

18. Hamann, E., Blevins, C., Franks, S. J., Jameel, M. I. & Anderson, J. T. Climate change alters plant–herbivore interactions. New Phytologist 229, 1894–1910 (2021).

19. Breiman, L. Random Forests. Machine Learning 45, 5–32 (2001).

20. Eyring, V. et al. Overview of the Coupled Model Intercomparison Project Phase 6 (CMIP6) experimental design and organization. Geoscientific Model Development 9, 1937–1958 (2016).

21. Grant, K., Kreyling, J., Dienstbach, L. F., Beierkuhnlein, C. & Jentsch, A. Water stress due to increased intra-annual precipitation variability reduced forage yield but raised forage quality of a temperate grassland. Agriculture, Ecosystems & Environment 186, 11–22 (2014).

22. Clauss, M. et al. The maximum attainable body size of herbivorous mammals: morphophysiological constraints on foregut, and adaptations of hindgut fermenters. Oecologia 136, 14–27 (2003).

23. Raghavan, G. V. Availability and use of shrubs and tree fodders in India. in Shrubs and tree fodders for farm animals: proceedings of a workshop in Denpasar, Indonesia, 24-29 July 1989 (IDRC, Ottawa, ON, CA, 1990).

24. Welti, E. A. R., Roeder, K. A., de Beurs, K. M., Joern, A. & Kaspari, M. Nutrient dilution and climate cycles underlie declines in a dominant insect herbivore. Proceedings of the National Academy of Sciences 117, 7271–7275 (2020).

25. Irob, K. et al. Browsing herbivores improve the state and functioning of savannas: A model assessment of alternative land-use strategies. Ecology and Evolution 12, e8715 (2022).

26. Berzaghi, F., Ben Abdallah, A., Ratshikombo, Z. & Bretagnolle, F. PNuts: Global dataset of plant nutritional values. bioRxiv (2022).

27. Abatzoglou, J. T., Dobrowski, S. Z., Parks, S. A. & Hegewisch, K. C. TerraClimate, a high-resolution global dataset of monthly climate and climatic water balance from 1958–2015. Scientific Data 5, 170191 (2018).

28. Fick, S. E. & Hijmans, R. J. WorldClim 2: new 1-km spatial resolution climate surfaces for global land areas. International Journal of Climatology 37, 4302–4315 (2017).

29. Harris, I., Osborn, T. J., Jones, P. & Lister, D. Version 4 of the CRU TS monthly high-resolution gridded multivariate climate dataset. Sci Data 7, 109 (2020).

30. Soil Survey Staff. Soil taxonomy: A basic system of soil classification for making and interpreting soil surveys. Agriculture handbook 436, 96–105 (1999).

31. Jung, M. et al. A global map of terrestrial habitat types. Scientific Data 7, 256 (2020).

32. Mariotti, F., Tomé, D. & Mirand, P. P. Converting Nitrogen into Protein—Beyond 6.25 and Jones’ Factors. Critical Reviews in Food Science and Nutrition 48, 177–184 (2008).

33. Bates, D., Mächler, M., Bolker, B. & Walker, S. Fitting Linear Mixed-Effects Models Using lme4. Journal of Statistical Software 67, 1–48 (2015).

