## Supplementary information for "Reduced nutritional quality of plants due to climate change"

The PDF file includes:

Materials and Methods

Extended Figs. S1 to S10

Extended Tables S1 to S2

References

Other Supplementary Materials for this manuscript include the following:

Extended Data S1

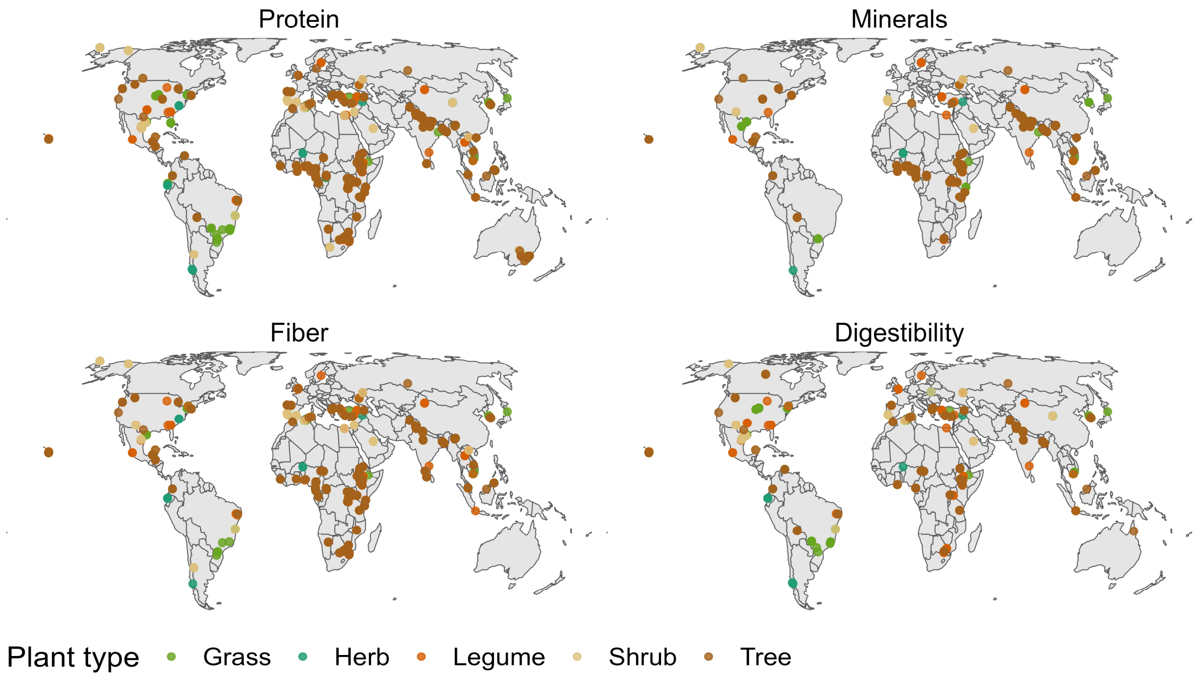


Fig. S1. Distribution of the nutritional properties data by plant type. Each point represents a location where one or more plant samples were collected in their natural growing environment.

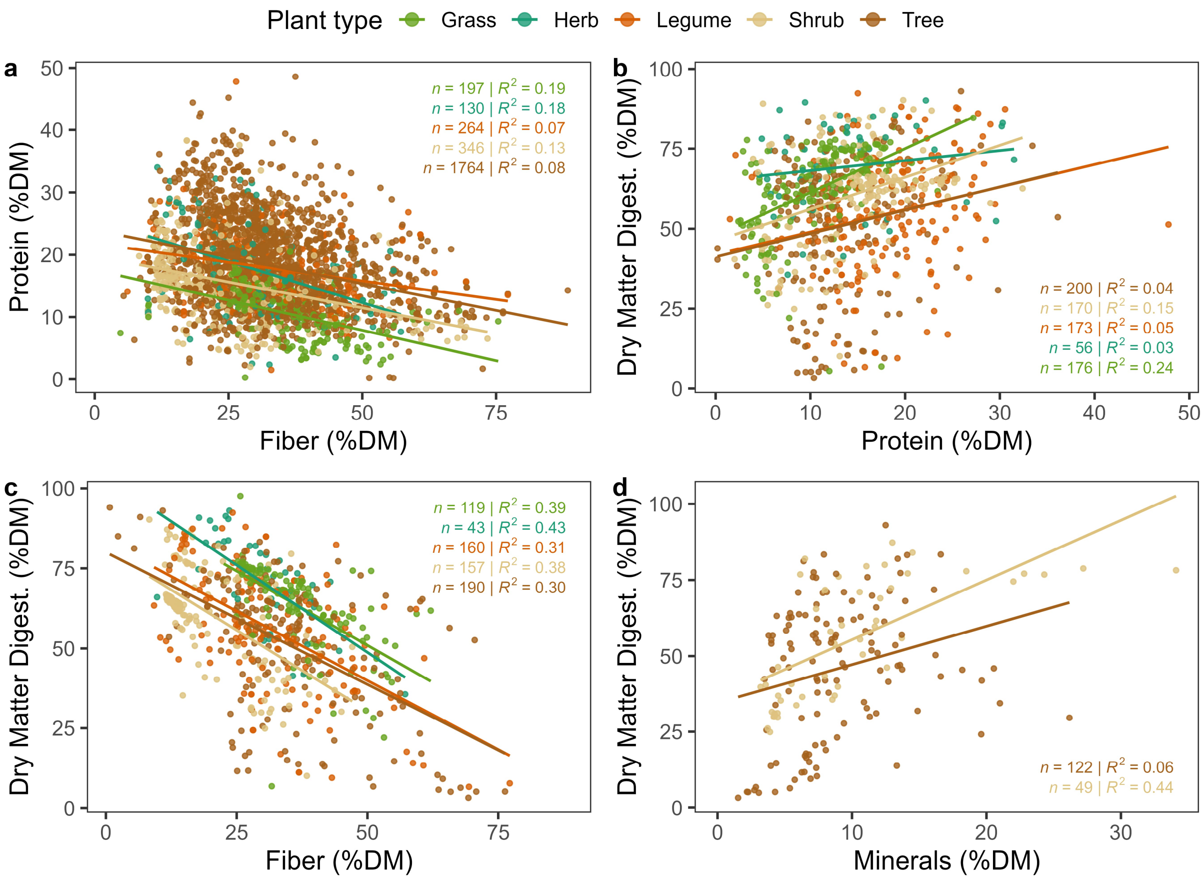


Fig. S2. Protein data as a function of fiber (a), and Crude protein, (b) fiber, and (c) minerals (d) as a function of dry matter digestibility. Only data from plant types with statistically-significant linear regressions are shown (p-value < 0.05). Complete test results are in the Extended Data s1.

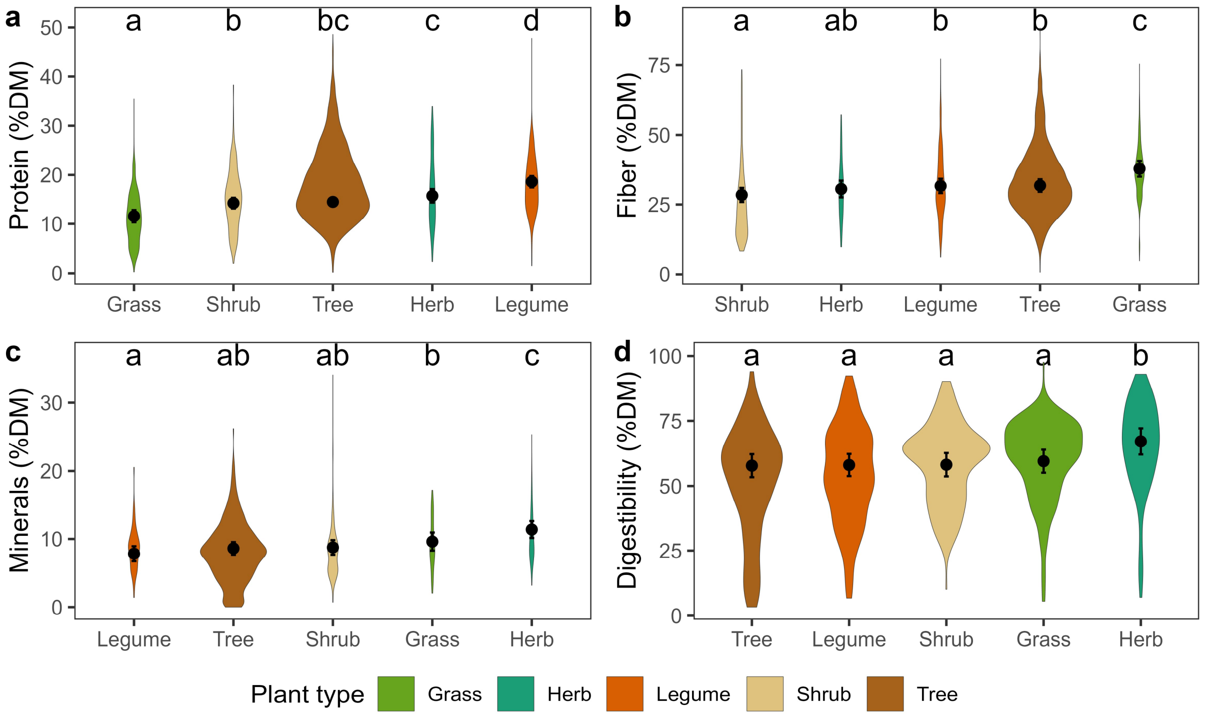


Fig. S3. Distribution of nutritional properties data across plant types. Plant types are ordered left to right in increasing order of mean (a) Protein, (b) fiber, (c) minerals, and (d) digestibility. Properties are all expressed as a % of dry matter. The width of violin plots is relative to the number of observations in each plot. Black dots and error bars represent estimated average ±95% confidence interval for each plant type. Means not sharing any letter are significantly different by the Sidak-test at the 5% confidence interval. Complete test results are in the Extended Data s1.

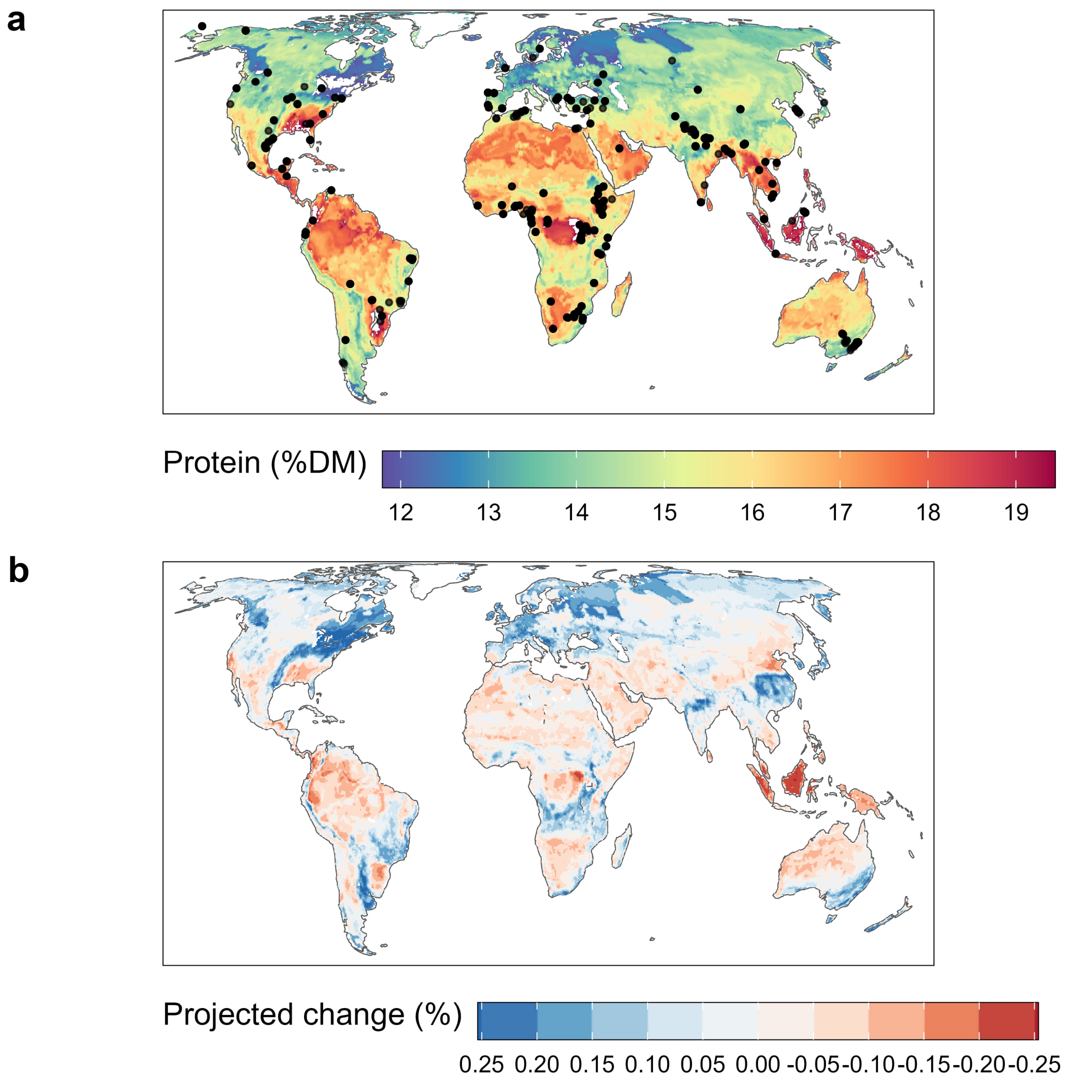


Fig. S4. Spatial distribution of plant protein in (a) present day and (b) projected relative changes in 2050 compared to present day. Each point in (a) represents a location where one or more plant samples were collected in their natural growing environment.

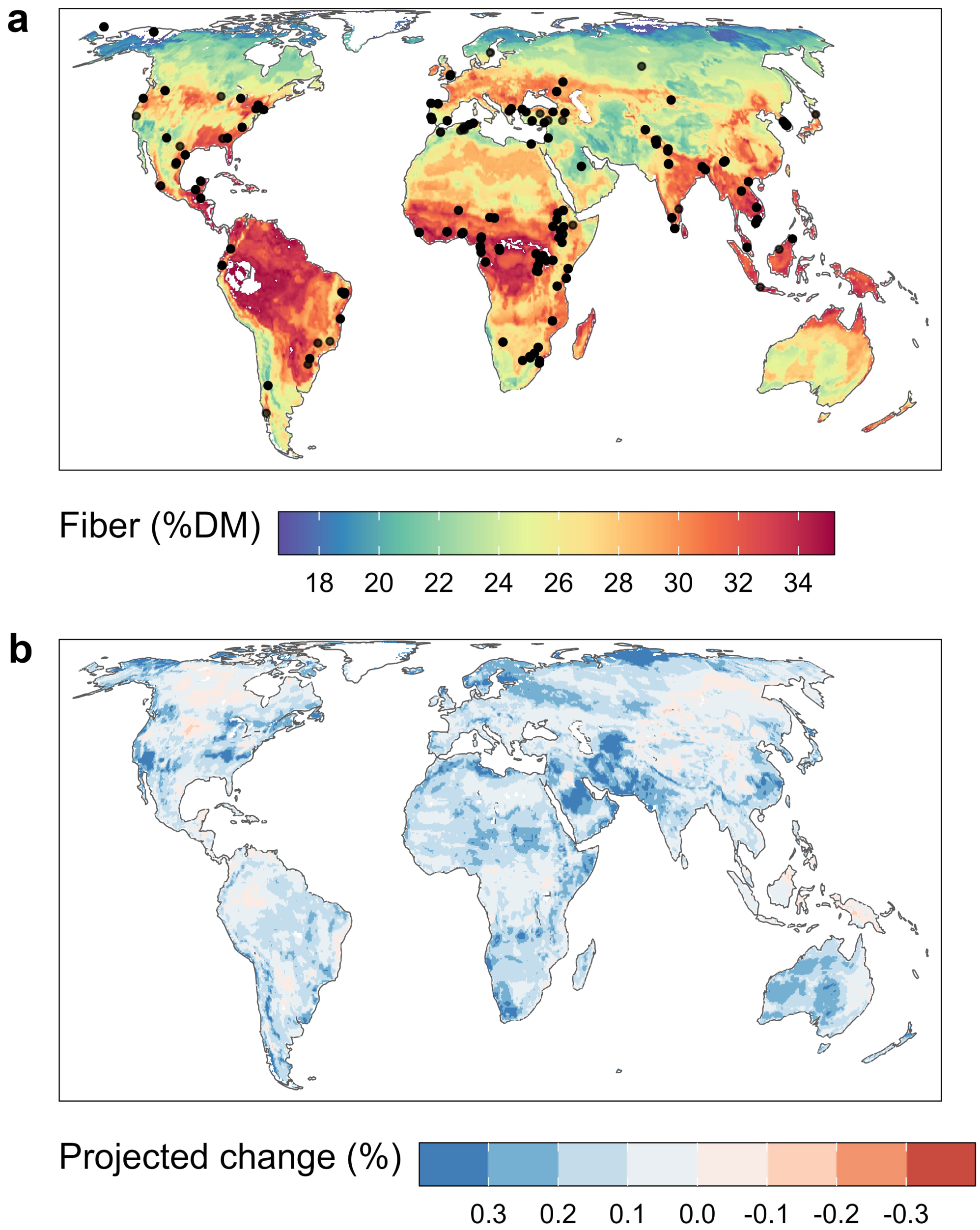


Fig. S5. Spatial distribution of plant fiber in (a) present day and (b) projected relative changes in 2050 compared to present day. Each point in (a) represents a location where one or more plant samples were collected in their natural growing environment.

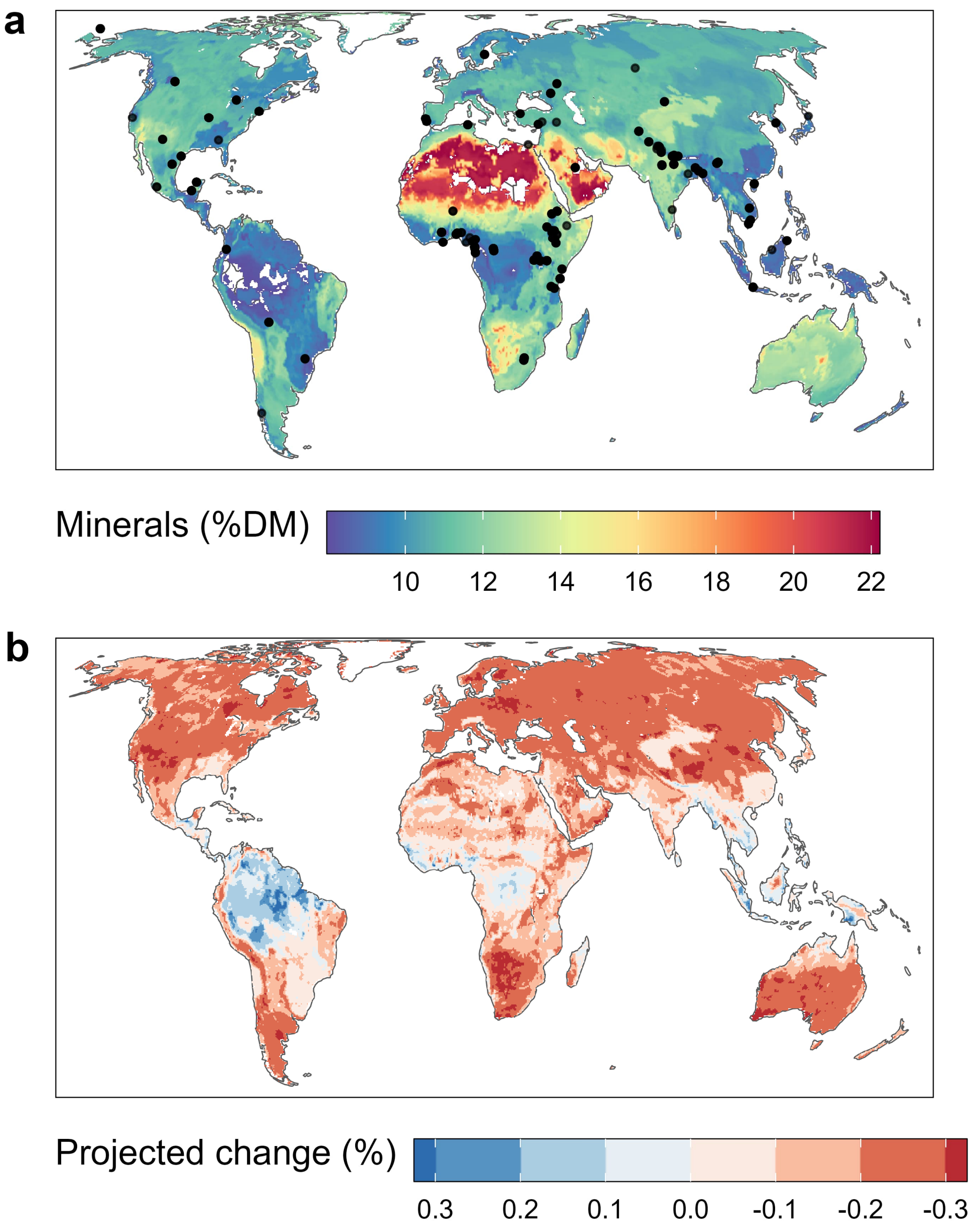


Fig. S6. Predicted spatial distribution of plant minerals in (a) present day and (b) projected relative changes in 2050 compared to present day. Each point in (a) represents a location where one or more plant samples were collected in their natural growing environment.

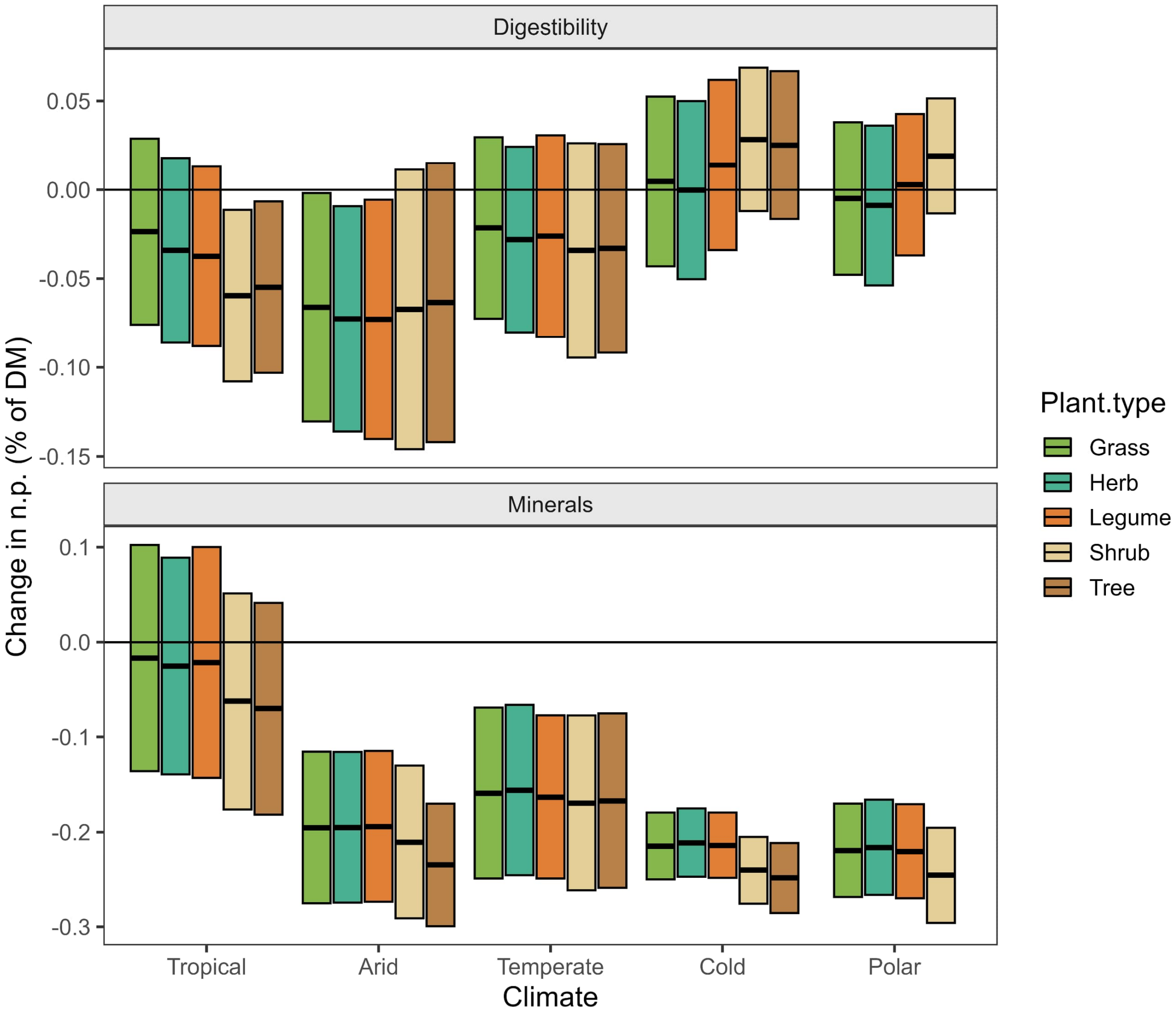


Fig. S7. Future changes in plants minerals and dry matter digestibility by 2050. The black horizontal centerlines indicate the average value for each bar plot. The top and bottom of the bars represent 1 ± standard deviation.

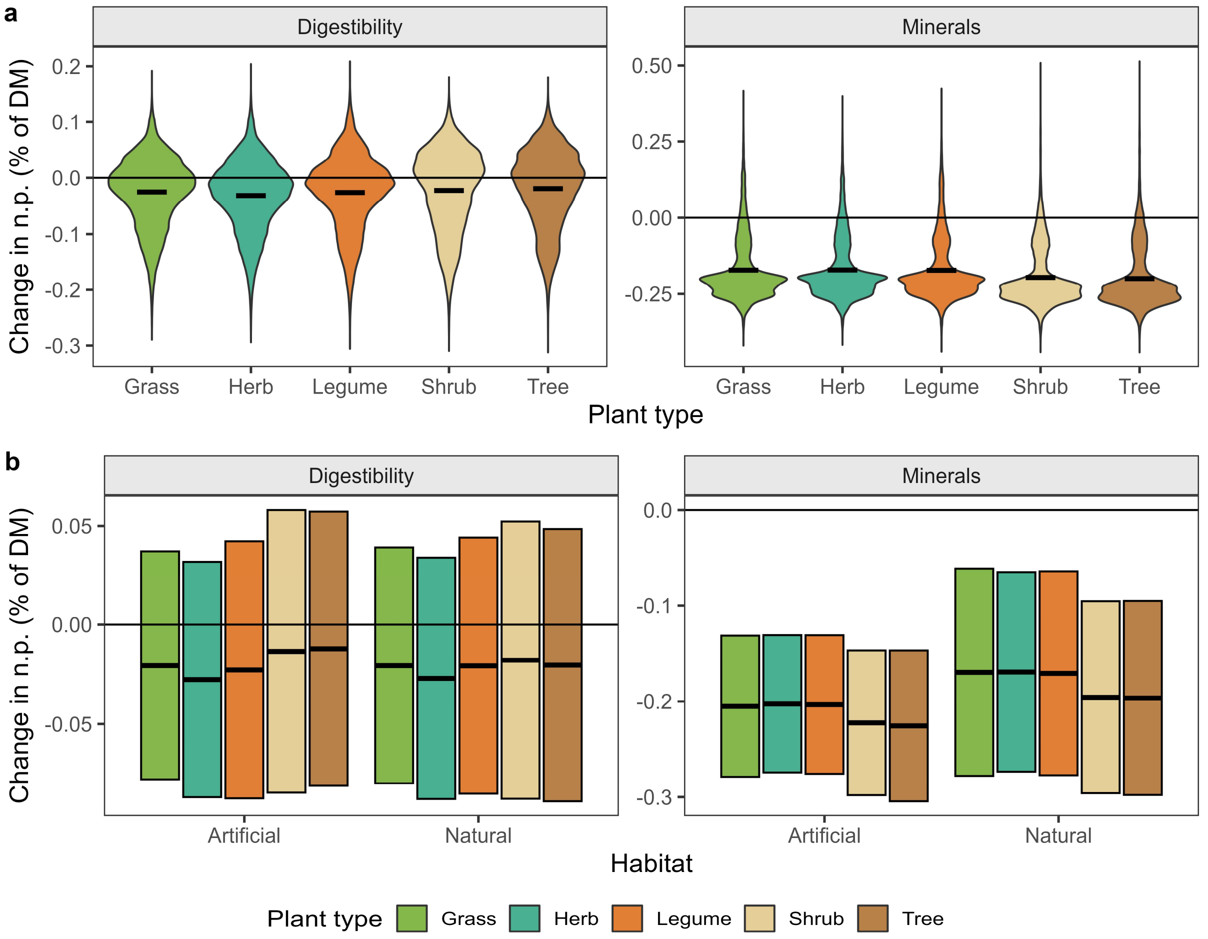


Fig. S8. Projected changes in plants minerals and dry matter digestibility by 2050 relative to present day. Analysis across (a) plant types and (b) habitat and plant types. The black horizontal centerlines indicate the average value for each bar or violin plot. The top and bottom of the bars represent 1 ± standard deviation. Please note the different y-scale in all plots.

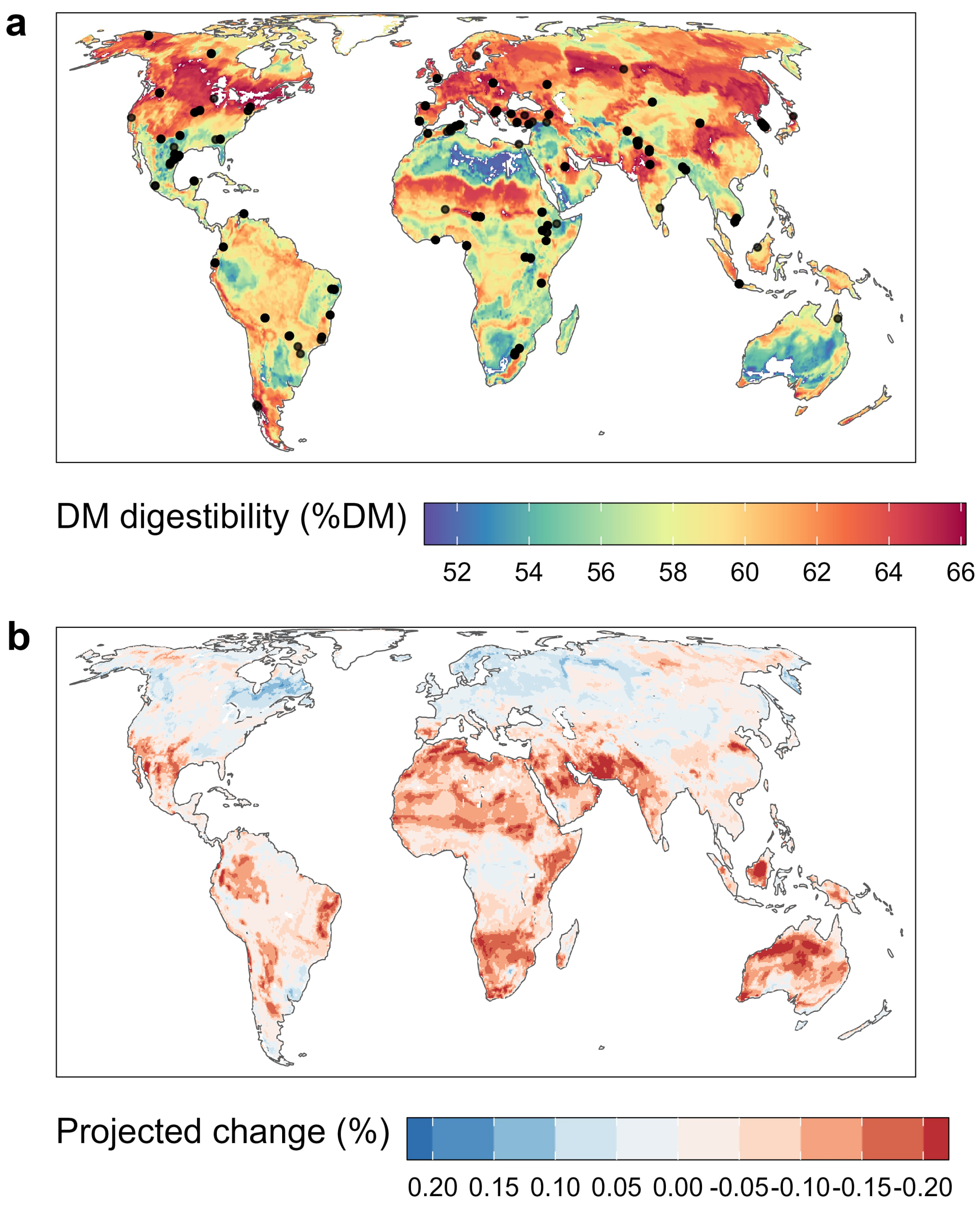


Fig. S9. Spatial distribution of plant digestibility in (a) present day and (b) projected relative changes in 2050 compared to present day. Each point in (a) represents a location where one or more plant samples were collected in their natural growing environment.

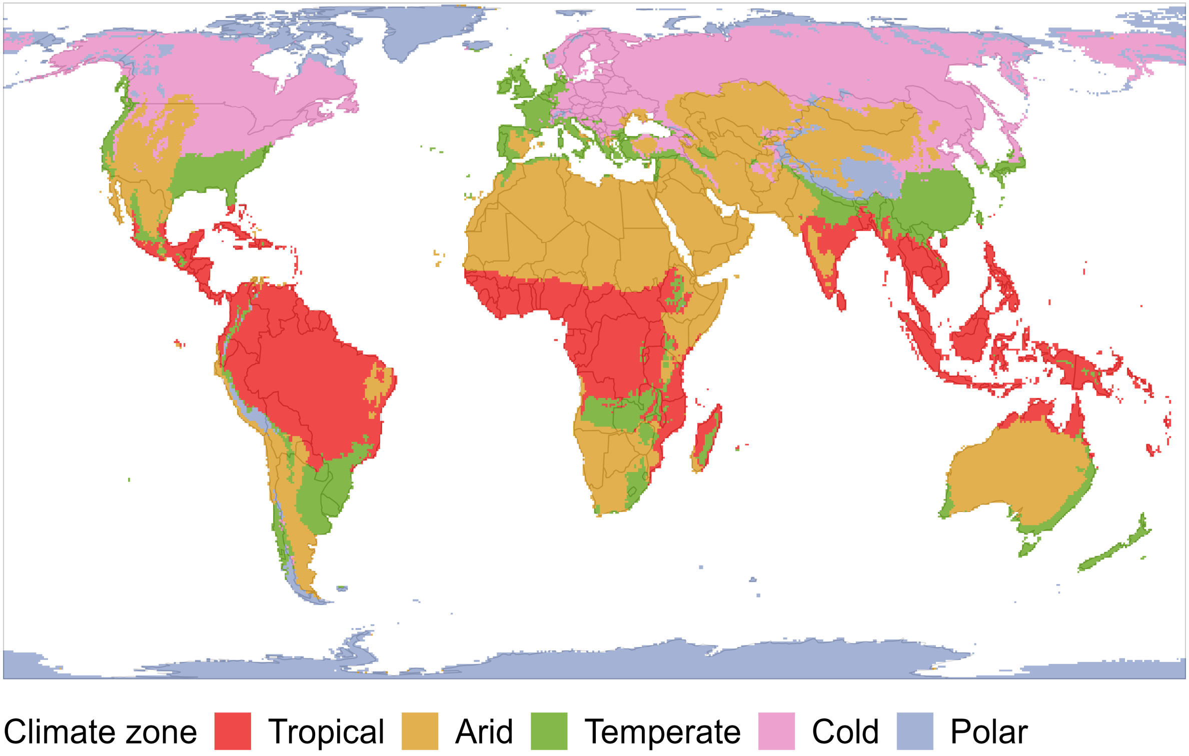


Fig. S10. Distribution of climate zones according to (*14*).

| Climate variable | Unit | 1968-2018 Source | Future scenarios source |
|---|---|---|---|
| Mean annual temperature | °C | CRU | WorldClim2/CMIP6 |
| Mean annual precipitation | mm/year | CRU | WorldClim2/CMIP6 |
| Precipitation seasonality | unitless index | CRU | WorldClim2/CMIP6 |
| Actual evapotranspiration | mm/month | TerraClimate | TerraClimate |
| Downward shortwave radiation | W/m² | TerraClimate | TerraClimate |
| $CO_2$ atmospheric concentration | ppm | NOAA/GML | CMIP6 |
| Soil type | categorical | USDA | - |

Table S1. Environmental variables sources and units used as predictors (in addition to plant type) in the random forest model.

| Property | % of variance explained | Root mean square error (% of DM) | Empirical data mean (% of DM) | Empirical data s.d. (% of DM) | Random forest mean (% of DM) | Random forest s.d. (% of DM) |
|---|---|---|---|---|---|---|
| Fiber | 54.9 | 9.54 | 32.5 | 12.5 | 25.8 | 4.6 |
| Minerals | 39.41 | 3.55 | 8.4 | 4.2 | 11.7 | 2.9 |
| Protein | 34.82 | 4.51 | 16.9 | 7.1 | 14.5 | 1.7 |
| Dry Matter Digestibility | 55.19 | 10.43 | 57.2 | 18.0 | 60.1 | 3.4 |

Table S2. Summary statistics of random forest model performance for the predicted nutritional properties. RMSE was computed using a test dataset.

**Data S1. (separate file)**

Results of statistical tests and complete list of changes in protein to fiber ratios of all countries.
